# Nearest-neighbor parameter for inosine-cytosine pairs through a combined experimental and computational approach

**DOI:** 10.1101/454124

**Authors:** Shun Sakuraba, Junichi Iwakiri, Michiaki Hamada, Tomoshi Kameda, Genichiro Tsuji, Yasuaki Kimura, Hiroshi Abe, Kiyoshi Asai

**Affiliations:** Graduate School of Frontier Sciences, The University of Tokyo, Japan; Quantum Beam Science Research Directorate, National Institutes for Quantum and Radiological Science and Technology, Japan; Faculty of Science and Engineering, Waseda University, Japan; Computational Bio Big-Data Open Innovation Laboratory (CBBD-OIL), National Institute of Advanced Industrial Science and Technology (AIST), Japan; Artificial Intelligence Research Center (AIRC), National Institute of Advanced Industrial Science and Technology (AIST), Japan; Department of Chemistry, Graduate School of Science, Nagoya University, Furo, Chikusa, Nagoya 464-8602, Japan; Division of Organic Chemistry, National Institute of Health Sciences, 3-25-26 Tonomachi, Kawasaki-ku, Kawasaki, Kanagawa 210-9501, Japan

**Keywords:** Inosine-cytosine pair, nearest-neighbor model, free energy, modified RNAs, molecular dynamics simulation

## Abstract

In RNA secondary structure prediction, nearest-neighbor parameters are used to determine the stability of a given structure. We derived the nearest-neighbor parameters for RNAs containing inosine-cytosine pairs. For parameter derivation, we developed a method that combines UV adsorption measurement experiments with free-energy calculations using molecular dynamics simulations. The method provides fast drop-in parameters for modified bases. Derived parameters were compared and found to be consistent with existing parameters for canonical RNAs. A duplex with an internal inosine-cytosine pair is 0.9 kcal/mol more unstable than the same duplex with an internal guanine-cytosine pair, and is as stable as the one with an internal adenine-uracil pair (only 0.1 kcal/mol more stable) on average.

## 1 Introduction

RNA molecules undergo various types of co‐ and post-transcriptional modifications. According to the MODOMICS database (Boccaletto et al., 2018), over 150 types of modifications are enzymatically introduced to transcripts across various species. Adenosine-to-inosine modification (a.k.a. A-to-I RNA editing) is catalyzed by adenosine deaminases acting on RNA (ADARs) converting adenosine in a RNA duplex (i.e., adenosine in A-U base-pairs) to inosine (Nishikura, 2010) and is one of the most prominent modifications in mammals. Currently, high-throughput sequencing methods, such as RNA-seq (Tan et al., 2017) and ICE-seq (Sakurai et al., 2014), provide the transcriptomic landscape of the A-to-I RNA editome. In the human transcriptome, more than 20,000 protein-coding/non-coding RNAs harboring 2.5 million A-to-I editing sites have been collected in the RADAR database (Ramaswami and Li, 2014).

Over the past decades, the secondary structures of RNA molecules have been experimentally investigated and computationally predicted, clarifying the relationships between their molecular functions and secondary structures (Hofacker, 2014; Seetin and Mathews, 2012). For example, pre-miRNAs and tRNAs are characterized by hairpin (Kozomara and Griffiths-Jones, 2014) and cloverleaf (Chan and Lowe, 2009) structures, respectively, despite the diversity of their primary sequences (Griffiths-Jones et al., 2003). Most of the computational predictions of RNA secondary structures are based on the nearest-neighbor (NN) model in which the stability of RNA duplexes is estimated as the summation of the free-energies of two adjacent base-pairs. These parameters, also known as NN parameters (Mathews et al., 1999; Xia et al., 1998), are derived from the thermodynamic measurement of RNA duplexes consisting of various pairs of the four standard RNA nucleotides (A, C, G, and U).

It is well known (Crick, 1966; Nishikura, 2010) that inosine forms a Watson-Crick type pair with cytosine and a wobble pair with uridine (Fig. 1A). However, computational predictions of secondary structures for inosine-containing RNA molecules are currently difficult because the NN parameters for inosine-cytosine (I-C) pairs have not been investigated, though the parameters for inosine-uridine (I*•*U) pairs have been reported (Wright et al., 2007). In an alternative method, inosines are treated as guanines in primary sequences (based on their similarity) for the purpose of predicting the secondary structures of inosine-containing RNAs. However, this method overestimates the stability of I-C pairs, because a G-C pair consists of three hydrogen bonds, whereas an I-C pair consists of only two.

**Figure 1:**
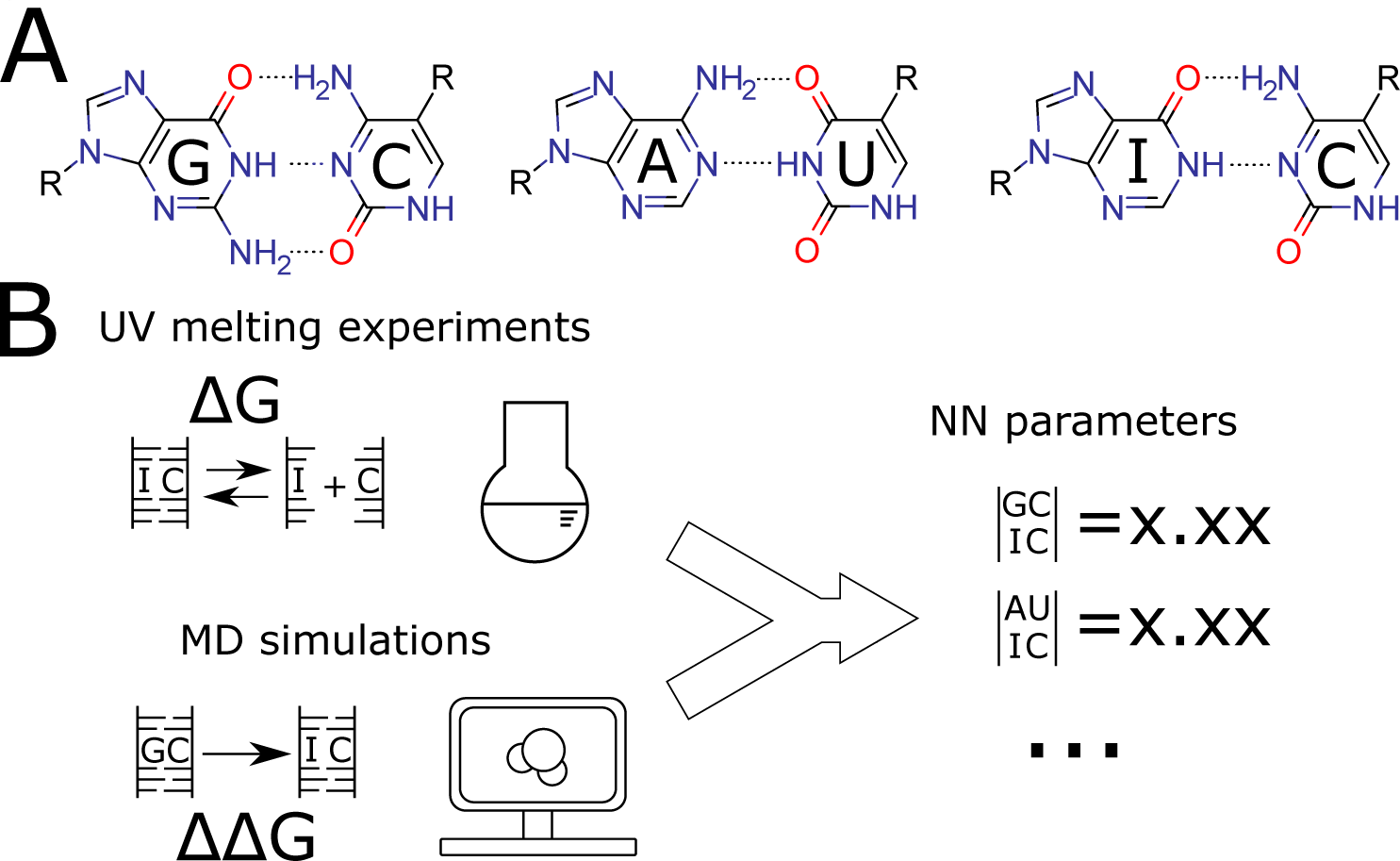
(A) Structure of the Watson-Crick pairs (G-C and A-U) and an I-C pair. (B) Schematic overview of the nearest-neighbor (NN) parameter derivation in this research. We combine the results from UV melting experiments with those from molecular dynamics simulations to obtain the NN parameters.

In previous research, we have shown that the combination of experimental free-energy measurements and free-energy calculations based on physicochemical simulation can be used to accurately estimate NN parameters (Nishida et al., 2018; Sakuraba et al., 2015). In the present work, we applied this method to determine the NN parameters for inosine-cytosine pairs (Fig. 1B). We compare the derived NN parameters with the corresponding parameters for other bases.

## 2 Results

### Experimental and computational duplex forming free-energy differences

Tables 1 and 2 list the experimentally observed 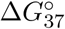 values and computationally derived 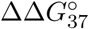 values, respectively. Each 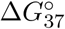 obtained from experimental measurement had high reproducibility, within a 0.17 kcal/mol error. Errors observed in computationally obtained 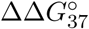 values were also within a similar range of the previously reported results.

**Table 1:**
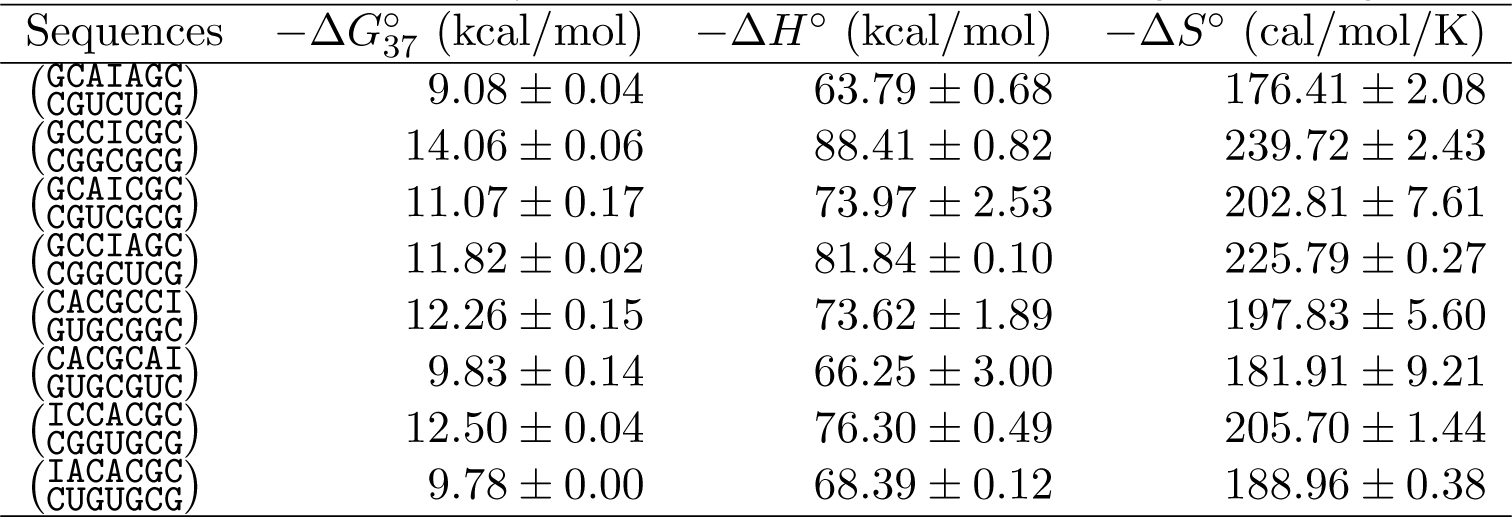
Experimentally measured RNA duplex-forming free energies.

**Table 2:**
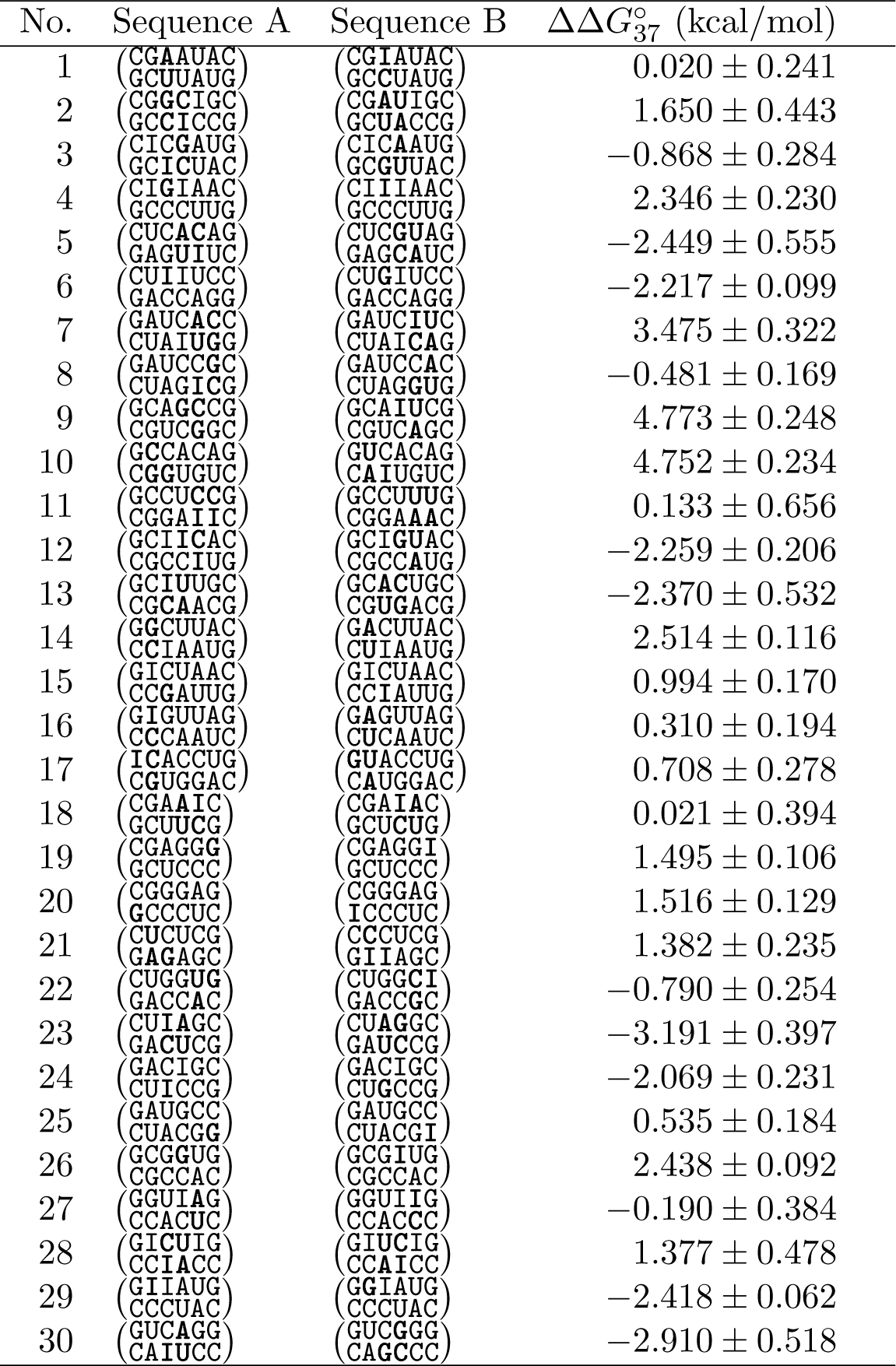
The relative difference in duplex-forming energy 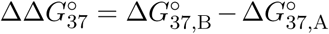 obtained from free-energy calculations.

### NN parameters and their validation

Combining both the experimentally observed and computationally derived free energies, we derived NN parameters for I*•*U pairs as Δ*G*_37,exp+comp_ (Table 3).

**Table 3:**
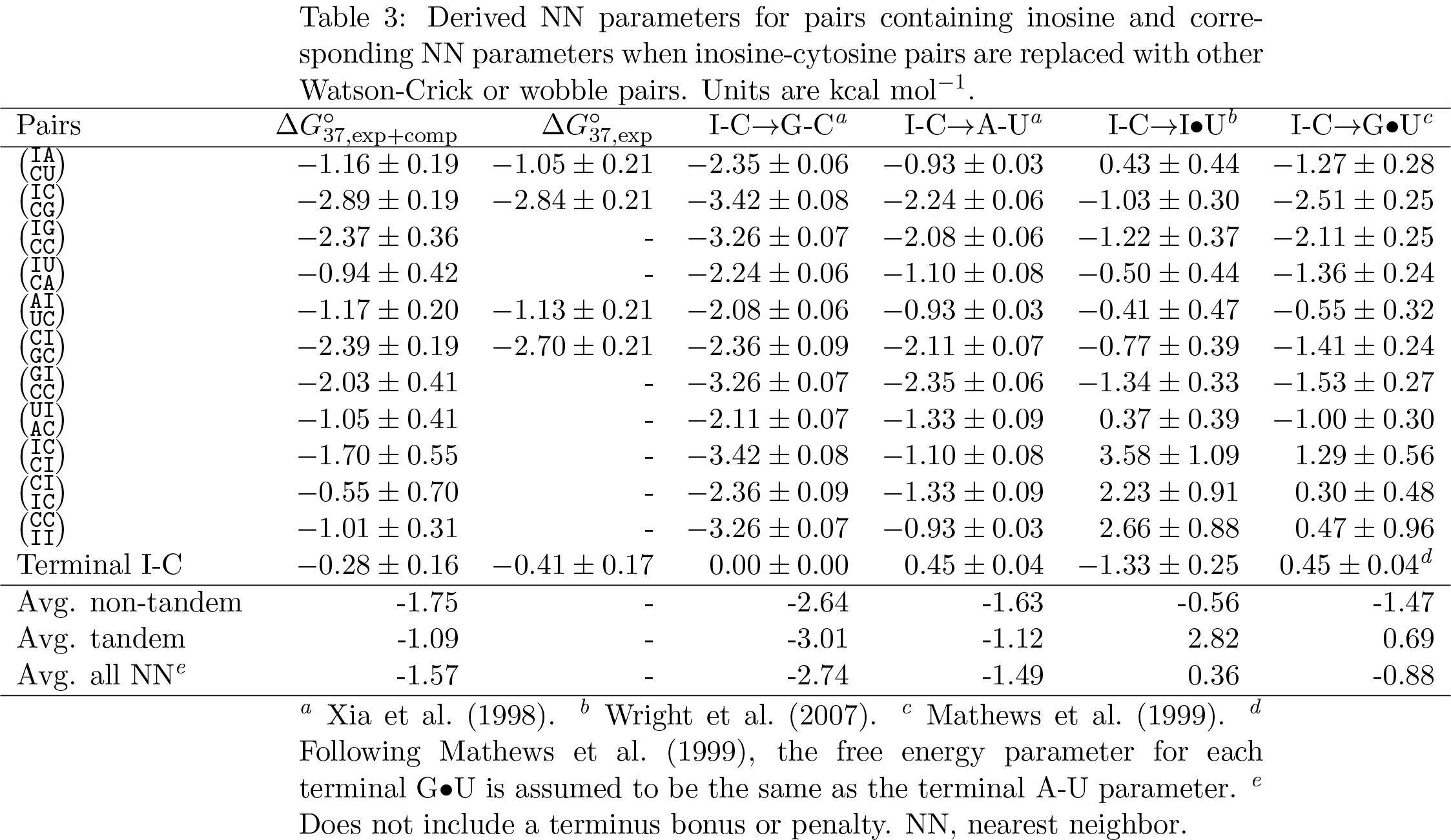
Derived NN parameters for pairs containing inosine and corresponding NN parameters when inosine-cytosine pairs are replaced with other Watson-Crick or wobble pairs. Units are kcal mol*^−1^*.

To validate these parameters, we first predicted the free-energy differences for the duplex formation based on the derived NN model and compared them with the experimental Δ*G* values and computational ΔΔ*G* values. Figure 2 presents the observed or calculated free energies versus the predicted free energies. As explained in the Methods section, the NN parameters were determined in favor of experimentally determined values. This is reflected in Fig. 2, as the experimental Δ*G* values have smaller deviations (mean absolute difference (MAD), 0.20 kcal/mol) from the predicted values while the calculated ΔΔ*G* values have larger deviations (MAD, 0.55 kcal/mol). Overall, the NN-predicted free energies are in good agreement with the observed Δ*G* values and calculated ΔΔ*G* values, demonstrating the consistency between the experimental Δ*G* values and the calculated Δ*G* values.

**Figure 2:**
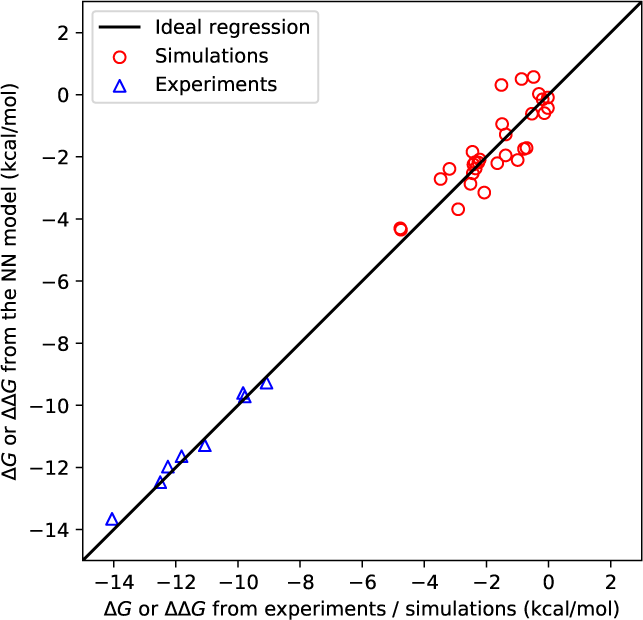
Comparison of nearest-neighbor (NN) model energies, and experimentally derived Δ*G* values or computationally derived ΔΔ*G* values. The sign of ΔΔ*G* values can be arbitrary inverted by swapping sequences A and B; we thus normalized the results to ensure ΔΔ*G* < 0.

The parameters were further validated using an off-by-one test. Because the parameters were determined with larger errors in the simulated free-energy difference, there is a risk that the results overfit the experimental free-energy difference. To address this possibility, we removed one experimental 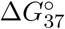 value from the dataset and derived the NN parameters. Then, we estimated the binding free-energy according to the NN model and compared it with the real experimental 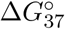 value. We repeated this procedure for all experimentally derived 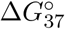 values (Fig. 3). Values from the model and the experiment agreed well (MAD = 0.41 kcal/mol and *R*^2^ = 0.94), indicating that the estimated free-energies are not sensitive to the elimination of a sample. We also tested whether the NN parameters are stable using an off-by-one analysis. The standard deviation of the parameters were within 0.12 kcal/mol, below the estimated standard deviations for any parameters. From these results, we concluded that the current parameter set is reasonably robust.

**Figure 3:**
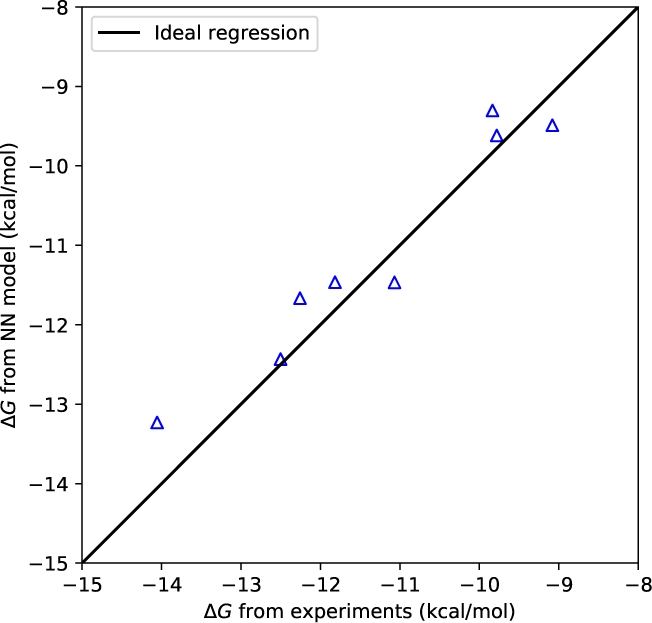
Comparison between the experimentally measured Δ*G* value and the prediction from a nearest-neighbor (NN) model that is determined by excluding a corresponding single experiment.

The NN parameters of all possible I-C and Watson-Crick pairs were all negative Δ*G* (favorable) values. For the case in which an inosine was at the 5′-end, the stabilities of the tandem pairs were ranked as follows: 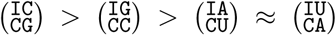. The difference between the parameters reflects the higher stability of guanine-cytosine (G-C) pairs compared with adenine-uracil (A-U) pairs, where the G-C pairs contain three hydrogen bonds between bases but the A-U pairs contain only two. The relative stability between the pairs with the same number of hydrogen bonds may reflect a more complex effect from the stacking. For the case in which an inosine was at the 3′-end, the stabilities of the tandem pairs were ranked as follows: 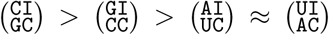. This again reflects the number of hydrogen bonds. For tandem inosines, the trend was as follows: 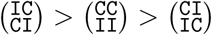.

We also compared the NN parameters with those obtained from experimental results alone (third column of Table 3). The differences between the parameters were marginal, all less than 0.40 kcal/mol. This is as expected because we used larger errors for the computationally obtained ΔΔ*G* values in the calculation of NN parameters.

## 3 Discussion

### Comparison to the related NN parameters

We compared the obtained NN parameters with the canonical NN parameters (Table 3). The fourth column (“I-C*→*G-C”) represents the NN parameters when I-C pairs were replaced by the G-C pairs (the column thus represents the NN parameters for 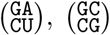, and so on). The NN parameters with I-C replaced by G-C (excluding any terminal bonus) had 1.17 kcal/mol lower free-energy differences on average compared with the I-C counterparts. The result is reasonable because each G-C pair has three hydrogen bonds, whereas each I-C pair has only two hydrogen bonds. The difference is thus considered to reflect the increased stability of G-C pairs with their additional hydrogen bond. The tendency is more prominent when parameters for only tandem I-C pairs are considered, where G-C containing pairs are 1.92 kcal/mol more stable than I-C counterparts. This is again reasonable because tandem G-C pairs have two additional hydrogen bonds compared with tandem I-C pairs. When I-C pairs were replaced by A-U pairs (“I-C*→*A-U”), the free-energy differences were comparable (0.08 kcal/mol in favor of I-C pairs). Again, this is considered to reflect the similar relative stability of A-U pairs (with two hydrogen bonds) and I-C pairs (also with two hydrogen bonds). The difference in the average parameters for tandem pairs was small as well (0.03 kcal/mol).

When the I-C pairs were replaced by I*•*U pairs, the free-energy difference of the duplex formation increased by 1.93 kcal/mol on average (Table 3, “I-C*→*I*•*U”). Even when the non-tandem NN parameters are considered, the NN parameters for I-C pairs are 1.19 kcal/mol more stable than the counterparts with I*•*U pairs. This result reflects the relative stability of Watson-Crick pairs and wobble pairs. Despite there being two hydrogen bonds between nucleotides in both pairs, wobble pairs are structurally disfavored compared with Watson-Crick pairs. The free-energy difference between tandem I-C pairs and tandem I*•*U pairs is 3.91 kcal/mol on average. This large difference indicates that stacking interactions of tandem wobble I*•*U pairs in RNA duplexes are strongly disfavored compared with tandem Watson-Crick I-C pairs. This tendency is also found in the comparison of NN parameters between tandem A-U (i.e., Watson-Crick) pairs (Xia et al., 1998) and tandem G*•*U (i.e., wobble) (Mathews et al., 1999) pairs. Tandem A-U pairs were generally more stable than tandem G*•*U pairs despite both having two hydrogen bonds between pairs (Table 3, “I-C*→*A-U” and “IC*→*G*•*U”), possibly due to the structural instability of consecutive wobble pairs.

### I-C bonus at RNA termini

In Table 3, 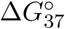 for a terminal I-C pair is −0.28 kcal/mol, indicating that there is a *bonus* when an I-C pair occurs at a terminus. This implies that I-C pairs at termini are more favored than G-C pairs. Indeed, the same tendency was also observed in the experimentally derived parameters (Table 3). Furthermore, terminal I*•*U wobble pairs have been reported (Wright et al., 2007) to be more stable than terminal G-C pairs (by −1.33 kcal/mol). Although the detailed mechanism is unclear, these results indicate that inosine-containing base pairs are favored on 5′and 3′-ends compared with C-G pairs. It is possible that single-stranded inosine is unstable because of structural restraints, entropic contributions, and/or solvation effects. Another possibility is that the observed stability of a terminal inosine reflects biases in the sequence design. Experimentally measured sequences containing I-C and I*•*U pairs are very limited in variation. All sequences with terminal I-C or I*•*U pairs in Table 1 and Wright et al. (2007) were GC rich (i.e., > 50% of base pairs are G-C). Errors that had accumulated due to this lack of variation may have caused this rather counterintuitive result. It has recently been shown that the accumulation of errors may sum up to unexpectedly large errors in the case of canonical NN parameters (Zuber et al., 2018). Limited variation in the sequences may have similar consequences in the case of both current I-C and I*•*U parameters.

### Future Research and Conclusions

Because the simulations inherently have a systematic bias from the experiments, the current NN parameters may be somewhat inaccurate. Ultimately, the parameters will be superseded by fully experimentally determined parameters. Meanwhile, the methodology of combining experiments with low-cost calculations should provide drop-in replacements for modified RNA parameters.

Recently, multiple RNA force fields(Aytenfisu et al., 2017; Kührová et al., 2018; Steinbrecher et al., 2012; Tan et al., 2018; Yang et al., 2017) have been proposed for the performance of precise RNA simulations. The main differences between the current RNA force field parameters and these new parameters are dihedral angle parameters(Aytenfisu et al., 2017) and van der Waals parameters (Steinbrecher et al., 2012; Tan et al., 2018; Yang et al., 2017). In the present research, because only the duplex-forming energy was considered, the difference in the dihedral angle parameters affects the stability of only single-stranded structures, whose contribution to the total free-energy difference may not be significant. The modification of the van der Waals parameters, on the other hand, may affect the stability of both single-stranded and duplex structures. These parameters may thus enable more accurate free-energy calculation and hence more accurate free-energy parameters. In parallel to these approaches, an implicit-solvent force field has recently been used to successfully estimate free energy (Chou et al., 2016). Combining these methodologies may contribute to the accurate, low-cost determination of NN parameters for various modified RNAs.

In the present research, only RNA duplex formation parameters were derived. Although parameters for the duplex formation are essential for RNA structure prediction, several other parameters are also known to contribute to more accurate RNA structure prediction. In particular, Zuber et al. (2017) recently reported that dangling and loop-initiation parameters strongly affect structural prediction accuracy for canonical RNAs. Combined with a recent report on the computational derivation of ΔΔ*G* of hairpins (Smith et al., 2018), methodologies for deriving non-stem parameters would be a valuable addition to the current parameter derviation scheme. Additionally, in the present research, the enthalpic and entropic terms were considered in only the experiments, not the simulations. Hence, the derived parameters cover Δ*G*s at only 37°C. Deriving parameters for a wider range of temperatures would enable prediction of structures at different temperatures. Additionally, recent research (Spasic et al., 2018) has suggested that using a multiple-state model improves the fit to UV-melting experiments. Adjusting the current methodology to fit such cases will be beneficial.

Finally, combined with the I*•*U wobble pair parameters by Wright et al. (2007), it is now possible to predict the secondary structure of inosine-containing sequences. The development of bioinformatic tools that enable structure prediction of inosine-containing sequences is important from a practical perspective.

In conclusion, we derived the NN parameters for RNAs containing I-C pairs by combining experiments and simulation. The derived parameters are consistent with the existing NN parameters for canonical bases. The current work also exemplifies a methodology for combining experiments with calculations in order to derive parameters for various modified RNAs.

## 4 Materials and Methods

### 4.1 NN model

In the NN model, the standard free-energy difference Δ*G^◦^* of forming a duplex is approximated by the sum of the energies for duplex initiation and those for two base pairs. For example, the standard free-energy difference for forming the duplex 5′-CAIA-3′ and 5′-UCUG-3′ at 37°C is approximated as

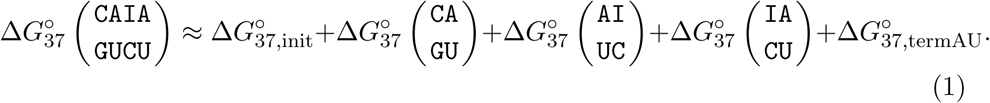

where 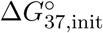 is the duplex initiation energy and 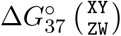 is the free-energy difference for each of the two neighboring base pairs. 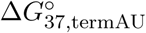 represents the penalty for each A-U pair appearing at the 5′and/or 3′ends. The free-energy difference for each two neighboring base pairs as well as penalties or bonuses at termini are represented by the NN parameters. The NN parameters are widely adapted as a fundamental parameter to estimate the secondary structure of RNAs (Hofacker, 2014; Seetin and Mathews, 2012). In this research, we only consider Watson-Crick type or wobble base pairs. Furthermore, unless otherwise stated, we only consider sequences at 37°C (i.e., 310K), and the temperature subscripts _37_ may be omitted from the notation.

As explained in the Introduction, our goal in this research is the derivation of the NN parameters for duplexes containing inosine-cytosine pairs. The NN parameters were typically determined from UV melting experiments; as also explained in the Introduction, this research has combined UV melting experiments with molecular simulations to estimate the parameters. Figure 1 represents the overview of approaches to determine the NN parameters. In addition to experimentally determined duplex-forming energies, we obtain the relative free-energy differences to supplement the parametrization.

### 4.2 UV adsorption measurements

The free-energy difference upon duplex formation for inosine-containing sequences were measured using UV adsorption experiments.

#### Sequence design

The sequences of inosine-containing RNAs were determined to meet the following criteria: (1) some NN parameters can be derived directly from experimental results, (2) self-pairing or staggered pairing is prevented, and (3) duplexes are stable enough to allow measurements around room temperature. We used DuplexFold in RNAStructure (Reuter and Mathews, 2010) to estimate the stability of RNA pairs, where we replaced inosine with guanine during the prediction. Table 1 lists the designed sequences.

#### Preparation of oligo RNAs

Solid phase synthesis of oligo RNAs was performed with an automatic DNA/RNA synthesizer (NTS M-2-MX, NIHON TECHNO Co., Ltd., Japan) using the following phosphoramidite monomers: A-TOM-CE Phosphoramidite, C-TOM-CE Phosphoramidite, G-TOM-CE Phosphoramidite, U-TOM-CE Phosphoramidite (10-3004, 10-3014, 10-3024, 10-3034, respectively; Glen Research, Sterling, VA), Inosine CED phosphoramidite (ANP-5680; ChemGenes, Wilmington, MA), and CPGs (Ac-C-RNA-CPG; 20-3315, Ac-G-RNA-CPG; 20-3324, Glen Research). For oligo RNAs with an inosine at the 3*^t^* end, Universal UnyLinker Support (2000°A, N-4000-20, ChemGenes) was used as a solid support. After solid phase synthesis, work-up procedures were performed as follows; the CPGs were washed with CH_3_CN, dried with flowing air, treated with a mixed solution of triethylamine and CH_3_CN (1:1, *v*/*v*), and kept at room temperature for 10 min. Then, the solution was removed and the CPGs were washed with CH_3_CN. Cleavage of the synthesized oligo RNAs from the CPGs and deprotection were performed with a mixed solution of concentrated NH_4_OH and 40% aqueous methylamine (1:1, *v*/*v*) at 35°C for 6 h or at room temperature for 12 h. The resulting RNA solution was filtered and transferred to another tube. This solution was concentrated to dryness using a centrifuge evaporator. The residue was treated with TBAF (1 M in THF), and the mixture was heated at 50°C for 10 min and vortexed. The mixture was then incubated at 35°C for > 6 h, and 1 M Tris-HCl buffer (pH 7.4) was added. Then, the volatiles were removed by a centrifuge evaporator. The resulting solution was desalted with a NAP-25 column (GE Healthcare, Chicago, IL). The collected fractions were concentrated with a centrifuge evaporator.

#### HPLC analysis and purification of oligo RNAs

HPLC purification and analysis were performed with a LaChrom Elite HPLC System (L-2455, L-2130, L-2485, HITACHI Co., Japan) equipped with a YMC HPLC column (Hydrosphere C18, 4.6 *×* 250 mm for analysis, 10 *×* 250 mm for purification, YMC Co., Ltd, Kyoto, Japan) with flow rates of 1 mL/min for analysis and 3 mL/min for purification. HPLC was conducted at room temperature and with a monitoring wavelength of 250 nm. The mobile phase consisted of solutions A (50 mM TEAA (pH 7.0) containing 5% CH_3_CN) and B (CH_3_CN), with a linear gradient of A/B = 100 : 0 to 70 : 30 used over 20 min. The collected fractions were lyophilized, and the resulting RNAs were re-dissolved in Milli-Q water (MilliporeSigma, Burlington, MA) and lyophilized by freeze drying once again.

#### MALDI-TOF MS analysis of oligo RNAs

The obtained RNA samples were characterized by MALDI-TOF MS (Ultraflex III, Bruker Daltonics, Billerica, MA) with a 3-hydroxypicolinic acid matrix. Calculated and observed MS values are summarized in Table S1.

*T_m_***measurements** The concentration of RNA stock solution (Milli-Q) was estimated by UV absorbance at 260 nm at room temperature using a NanoDrop 2000 (Thermo Fisher Scientific, Waltham, MA USA). The molar extinction coefficient of oligo RNAs was calculated using Oligo Analyzer 3.1 (Integrated DNA Technologies, Coralville, IA), an online software tool. Equal moles of complementary single strands of RNA were mixed to form the duplex. This solution was lyophilized, and then the RNAs were redissolved in 20 mM sodium cacodylate, 1 M NaCl, and 0.5 mM Na_2_EDTA (pH 7.0). The *T_m_* values of the samples were recorded on a spectrophotometer (V-650; JASCO, Tokyo, Japan) equipped with an 8-microcell changer and Peltier temperature controller system using a heating rate of 2*°*C/min from 10°C to 90*°*C to obtain curves of absorbance at 260 nm versus temperature. Spectrum manager software (JASCO) was used to fit melting curves to a two-state model to calculate *T_m_* values. *T_m_* values at different concentrations (2, 4, 6, 8, 10, 50, 100, 120, and 150 *µ*M) were used to calculate thermodynamic parameters.

#### Data analysis

The duplex-forming free-energy difference was estimated from the regression between the concentration of the RNA (*C_t_*) and the melting temperature (*T_m_*). We followed the procedure detailed by Xia et al. (1998) to derive 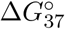, Δ*H^°^*, and Δ*S^°^* upon the duplex formation. We calculated the parameters from three independent measurements. Averages and the standard deviations are listed in Table 1.

### 4.3 Free-energy calculation

Although the direct calculation of free-energy differences upon duplex formation is possible (Deng and Roux, 2009; Tan et al., 2018; Wang et al., 2006), such calculation often requires considerable computational resources and/or dedicated hardware to achieve an accuracy comparable to that based on experiments. This is because there are large structural changes between the two end states (i.e., the duplex form and the unbound form), and the various conformational changes from two states should be sampled to achieve better convergence. Instead, we considered the calculation of the *relative* free-energy difference. Here, we considered the relative freeenergy difference with respect to the mutations to nucleotide bases, that is, the *difference* in duplex-forming energies between two RNA duplexes ΔΔ*G*_AB_ = Δ*G*_B_ *−* Δ*G*_A_. This kind of free-energy difference causes smaller conformational changes, and thus the resulting relative free-energy difference converges faster (Krepl et al., 2013; Sakuraba et al., 2015). Similar approaches have also been demonstrated to be computationally efficient in analyses of protein-drug complexes (Wang et al., 2015). Furthermore, we have shown that such relative free-energy differences can also be used to estimate NN parameters in combination with experiments (Nishida et al., 2018). The relative binding free-energy changes were calculated for various inosine-containing RNA duplexes using the free-energy perturbation method.

#### Sequence design

We chose pairs of inosine-containing duplexes so as to meet the following goals: (a) the pairs cover a variety of the NN parameters containing inosine, (b) the mutations between duplexes do not contain unnecessary canonical NN changes, (c) all duplexes are stable enough to hold a duplex form at 37°C, (d) the sequence does not contain excessive repetition, (e) there are only two or fewer base pair mutations, (f) termini are paired as G-C or I-C (for stability), and (g) sequence lengths are 6 or 7 bases. We optimized the pairs using in-house software to meet the criteria above. During the design process, we used the Vienna RNA package (Lorenz et al., 2011) to predict the stability of various duplexes, with inosine bases replaced by guanine bases during the prediction.

#### System setup

The relative free-energy differences were calculated with the free-energy perturbation method. We used the same protocol as described by Sakuraba et al. (2015) to calculate the free-energy difference; for the detailed procedure, please consult this previous study. Parameters associated with atoms (van der Waals radii and charges) were gradually changed between two end states (pairs A and B) to compute the free-energy difference. The 1-1-48 soft-core potential function (Pham and Shirts, 2011) was used to remove numerical instability during the calculation. Both charges and the van der Waals radii were treated with the soft-core potential.

For the 30 duplex pairs listed in Table 2, all-atom, explicit-water models were constructed. The AMBER14SB (AMBER99 + bsc0 + *χ*OL3 (Cheatham III et al., 1999; Perez et al., 2007; Zgarbová et al., 2011) for nucleic acids) force field and a monovalent cation force field developed by Joung and Cheatham (Joung and Cheatham, 2008) were used to model the system. Water molecules were modeled using TIP3P, and the system was solvated with 1 M NaCl solution. The atomic charges of an inosine molecule were determined by the RESP method (Bayly et al., 1993; Cieplak et al., 1995). The dihedral angle parameter with respect to the *χ* angle was set equal to that from the adenine molecule in a *χ*OL3 force field (Zgarbová et al., 2011). The total number of atoms in the system was between 13,741 and 16,481. For each pair of duplexes, the free-energy difference with respect to the mutation was calculated using the weighted histogram analysis method (multistate

Bennett acceptance ratio method) (Kumar et al., 1993; Shirts and Chodera, 2008; Souaille and Roux, 2001). The Hamiltonian replica exchange method (Fukunishi et al., 2002) was used to sample various conformations between two end states (28 replicas in total). Free-energy differences were calculated for the duplex state and two single-stranded states, and the total relative free energy difference was calculated according to the thermodynamic cycle in Fig. 4. For each state, the production run was performed for 20 ns (20 ns *×* 28 replicas *×* 3 states *≈* 1.7 *µ*s for each pair of duplexes). For the duplex states of pairs 17, 19, 20, 22, and 25, we restrained the termini with I-C (and corresponding termini at the other state) to keep Watson-Crick type pairs because inosine bases at termini are unstable at 37°C and often result in a dissociation between the two strands. The restraint was realized by applying a potential that includes a penalty if the distance between N1 of inosine and N3 of cytosine exceeds 4 °A, with a force constant of 10^4^kJ/mol/nm^2^.

**Figure 4:**
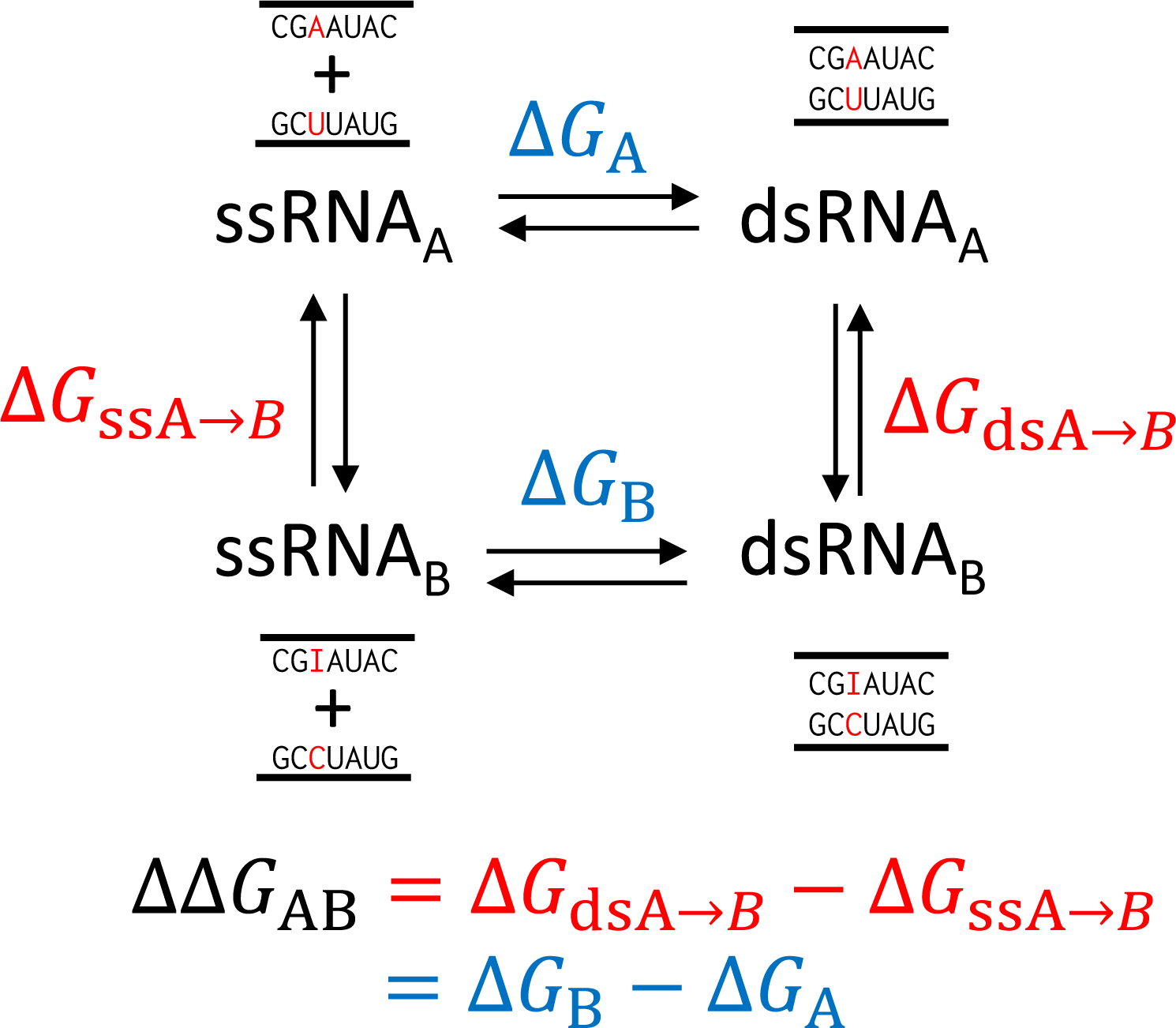
The thermodynamic cycle considered in this research. From UV melting experiment Δ*G* values were directly measured, while from the molecular dynamics simulations, Δ*G* values corresponding to the mutation of *A* → *B* were calculated as ΔΔ*G*_AB_.

### 4.4 NN parameter estimation

We derived the NN parameters using both the experimentally derived freeenergy differences and the computationally derived relative free-energy differences. We used a method described by Nishida et al. (2018) to combine experimental and computational results. Here, we briefly describe the method to derive the NN parameters.

#### Linear least square model

With the NN model, the RNA duplex-forming energy can be approximated as the matrix-vector product

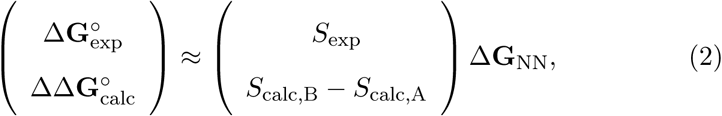

where 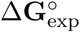 is a vector representing experimentally measured Δ*G^°^* values, 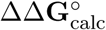 is a vector representing calculated ΔΔ*G^°^* values, and Δ**G**_NN_ is a vector representing the NN parameters we wish to obtain. *S*_exp_ and *S*_calc_ are matrices constructed as follows: each row of *S*exp represents the number of occurrences of NNs in the duplex, whereas each row of *S*_calc,A_ and *S*_calc,B_ represents the number of occurrences in sequences before and after the mutation, respectively. In this research, we only consider the NN parameters containing I/C pairs. This can be achieved by subtracting offset energies based on the occurrences of all the canonical NN pairs from 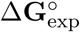 and 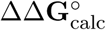.

From Eq. (2), the NN parameters are determined by solving the least square fitting problem. The parameters are readily obtained as

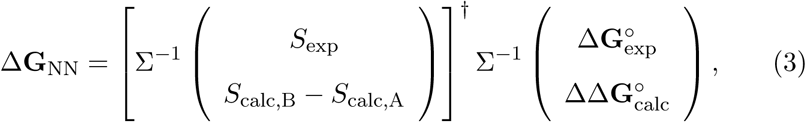

where *M^†^* is the Moore-Penrose pseudoinverse of the matrix *M*, and Σ is a diagonal matrix whose diagonal elements are standard deviations of all experiments and calculations combined. In this work, we assumed that all experiments and all calculations are independent and that their errors are not correlated. Expected standard deviations can be derived similarly(Nishida et al., 2018) and are presented alongside the parameters in Table 3.

#### Derivation of parameters

The simulation inherently has a *systematic* bias compared with the experimental counterpart. Although both the simulation and experiments have biases compared with the “true” values of Gibbs free energy (or differences in Gibbs free energy), the bias from the simulation is considered to be much larger than that from the UV melting experiment. Consistent with the case of the canonical NN parameter, in this research, we consider that the values obtained from the UV melting experiments represent the “true” values of the Gibbs free energy. We thus consider that the values from UV experiments have only statistical errors, while the values from simulations have both statistical errors and systematic biases. In our previous work (Sakuraba et al., 2015), we observed a mean absolute deviation of 0.55 kcal/mol in the simulation of canonical bases. Based on this result, to compensate for the systematic biases, we offset the estimated statistical errors from the free-energy simulation by 0.50 kcal/mol. This protocol adjusts the NN parameters to favor experimentally measured free energies, reflecting the uncertainty arising from the systematic bias in the computationally derived free energies. For a comparison, we also derived the NN parameters from the simulation-derived free energies only. Derived parameters are presented in Table SII. Derived parameters indicated a slight deviation from the experimental values as presented in Fig. SI, therefore, we concluded that the inclusion of experimental parameters is necessary for the accurate parameter derivation.

## Acknowledgement

The computations were performed using ACCMS, Kyoto University, Japan, and the Research Center for Computational Science, Okazaki, Japan. This work was supported by JSPS KAKENHI Grant Numbers 16H02484 to SS, MH, TK, HA, and KA, 16K17778 to SS, and 16K16143 to JI, and by a MEXT Grant-in-Aid for Scientific Research on Innovative Areas (16H06279) to KA.

## Supporting Information

Characterization data for the synthesized inosine oligo RNAs, NN parameters from only simulations, and parameter files we used for inosine simulation are available in the supporting information.

## List of Figures

1. A) Structure of the Watson-Crick pairs (G-C and A-U) and an I-C pair. (B) Schematic overview of the nearest-neighbor (NN) parameter derivation in this research. We combine the results from UV melting experiments with those from molecular dynamics simulations to obtain the NN parameters. 28
2. Comparison of nearest-neighbor (NN) model energies, and experimentally derived Δ*G* values or computationally derived ΔΔ*G* values. The sign of ΔΔ*G* values can be arbitrary inverted by swapping sequences A and B; we thus normalized the results to ensure ΔΔ*G <* 0. 29
3. Comparison between the experimentally measured Δ*G* value and the prediction from a nearest-neighbor (NN) model that is determined by excluding a corresponding single experiment. 30
4. The thermodynamic cycle considered in this research. From UV melting experiment Δ*G* values were directly measured, while from the molecular dynamics simulations, Δ*G* values corresponding to the mutation of *A* → *B* were calculated as ΔΔ*G*_AB_ 31

## List of Tables

1. Experimentally measured RNA duplex-forming free energies. 33
2. The relative difference in duplex-forming energy 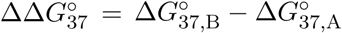 obtained from free-energy calculations. 34
3. Derived NN parameters for pairs containing inosine and corresponding NN parameters when inosine-cytosine pairs are replaced with other Watson-Crick or wobble pairs. Units are kcal mol*^−1^* 35

